# Unveiling reversibility and plasticity in cardiac hypertrophy: insights from a transverse aortic constriction-release model

**DOI:** 10.1101/2024.07.07.602358

**Authors:** Manabu Shiraishi

**Affiliations:** Department of Cardiovascular Surgery, Saitama Medical Center, Jichi Medical University, Saitama, Japan

**Keywords:** Heart failure, Hypertrophy, Fibrosis, Angiogenesis, Transverse aortic constriction, Reversibility

## Abstract

Transverse aortic constriction (TAC) is a well-established animal model used to study the pathomechanisms of pressure overload-induced heart failure. A number of studies have shown that treatment of the heart failure in this model may reverse the associated hypertrophy and fibrosis. However, because no TAC-release model in which hemodynamics improve upon alleviation of the physical stenosis has yet been established, the histologic changes and regulatory molecular biological mechanisms underlying the reversibility of cardiac hypertrophy and fibrosis are unknown. This study was conducted to establish an animal TAC-release model and thereby investigate the mechanisms that govern reversibility and plasticity of myocardial hypertrophy, fibrosis, and angiogenesis. TAC surgery was performed on rats, and 4 weeks later TAC release was achieved by cutting the constricting threads. TAC-subjected heart exhibited severe myocardial hypertrophy, fibrosis, and increased angiogenesis, along with diastolic dysfunction. Heart released from TAC showed reduced hypertrophy and fibrosis and improved diastolic function. Gene expression analysis uncovered regulator of calcineurin 1 (*Rcan1*) as a key player in cardiac function and histologic changes after TAC release. *Rcan1* knockdown exacerbated myocardial hypertrophy and fibrosis in heart released from TAC. The left ventricular afterload relief model revealed that increased oxidative stress and *Rcan1* upregulation, which suppresses the calcineurin-NFAT pathway, are key to structural and functional recovery from pressure overload-induced cardiac hypertrophy.

## Introduction

Left ventricular pressure overload, as represented by systemic hypertension and aortic stenosis, induces progressive fibrosis, markedly decreasing myocardial compliance, often leading to lethal outcomes^1^. Animal models, such as transverse aortic constriction (TAC), have been developed to study the pathophysiology of pressure overload-induced heart failure, with numerous experiments exploring the underlying cellular and molecular mechanisms^2–4^. Elevated pressure in the left ventricle triggers the binding of stress hormones, especially angiotensin II and noradrenaline, to myocardial cell surface receptors, activating intracellular signaling pathways. This culminates in increased expression of transcription and cell growth factors, inducing myocardial hypertrophy. However, interventions can block these pathways, inhibiting cardiomyocyte apoptosis and impeding progression of the hypertrophy, thereby regulating cardiac reversibility and plasticity^5, 6^. Myocardial fibrosis resulting from left ventricular pressure overload impairs heart function by excessive accumulation of extracellular matrix (ECM) produced by activated myocardial fibroblasts^7, 8^. Extensive fibrosis is generally irreversible, but studies have suggested the potential for reversal with treatment addressing underlying diseases in experimental models^9, 10^. Cardiac hypertrophy has also been linked to accelerated angiogenesis^11^, which occurs to support increased myocyte proliferation and metabolism. Although the mechanisms of reversibility of angiogenesis have not been fully clarified, it appears that reducing cardiac overload and hypertrophy through appropriate treatment reverses angiogenesis by apoptosis-induced elimination of vascular cells and by decreasing growth factors, reducing inflammation and angiogenic demand^12^. Understanding the mechanisms responsible for reversibility and plasticity of myocardial hypertrophy, fibrosis, and angiogenesis is crucial for the development of heart failure therapies. However, understanding of the regulatory mechanisms involved in reversal of cardiac hypertrophy, fibrosis, and angiogenesis following hemodynamic changes remains incomplete because there is no animal model in which left ventricular pressure load resulting from the physical stenosis introduced by TAC is decreased. To bridge this gap in our understanding, a TAC-release model was created, allowing for comprehensive profiling of the structure, function, and histology of the heart. The goal was to elucidate the molecular mechanisms underlying reversibility and plasticity particular to myocardial hypertrophy, fibrosis, and angiogenesis.

## Results

### Characteristics of heart released from TAC

To elucidate the mechanism by which myocardial homeostasis is maintained in myocardial environments with under different hemodynamic loads, rats were divided into groups for establishment of the following conditions: a normal heart (NH) condition (for control), a pressure-overload heart condition, with hearts by subjected to TAC (TAC-subjected heart), and an overload relief condition, with removal of the constricting threads 4 weeks after TAC surgery (heart released from TAC), as previously described^13^ (Fig. 1a). M-mode echocardiography revealed a significant increase in left ventricular posterior wall thickness at diastole (LVPWd), reduction in left ventricular dimension at diastole and at systole (LVDd and LVDs), and volume of TAC-subjected heart compared to measures in normal heart. These alterations improved in heart released from TAC, seemingly due to a reduction in left ventricular afterload. Left ventricular ejection fraction (LVEF) and other indicators of left ventricular contractility did not differ significantly between conditions, presumably due to compensatory left ventricular hypertrophy (Fig. 1b). The causes of these echocardiographic changes were then investigated in terms of histology. A significant increase in the cross-sectional area around the short-axis of individual cardiomyocytes (viewed from the transverse section) was observed in TAC-subjected heart compared to that area in normal heart, and in heart released from TAC, the area decreased to the size observed in normal heart (Fig. 1c). The amount of collagen deposition was significantly increased in the TAC-subjected heart versus that in normal heart, and this increase was attenuated in heart released from TAC (Fig. 1d). Capillary density was increased in TAC-subjected heart and decreased in heart released from TAC (Fig. 1e). The data suggest that hypertrophy of individual cardiomyocytes and increased collagen deposition in the interstitium enhance left ventricular wall thickness and that angiogenesis is augmented by the increased oxygen demand of the hypertrophied myocardium. Furthermore, the reduction in left ventricular afterload effected by TAC release suppressed cardiac hypertrophy, fibrosis, and angiogenesis. Interestingly, reversible histologic changes were observed in response to the hemodynamic changes.

**Figure 1.**
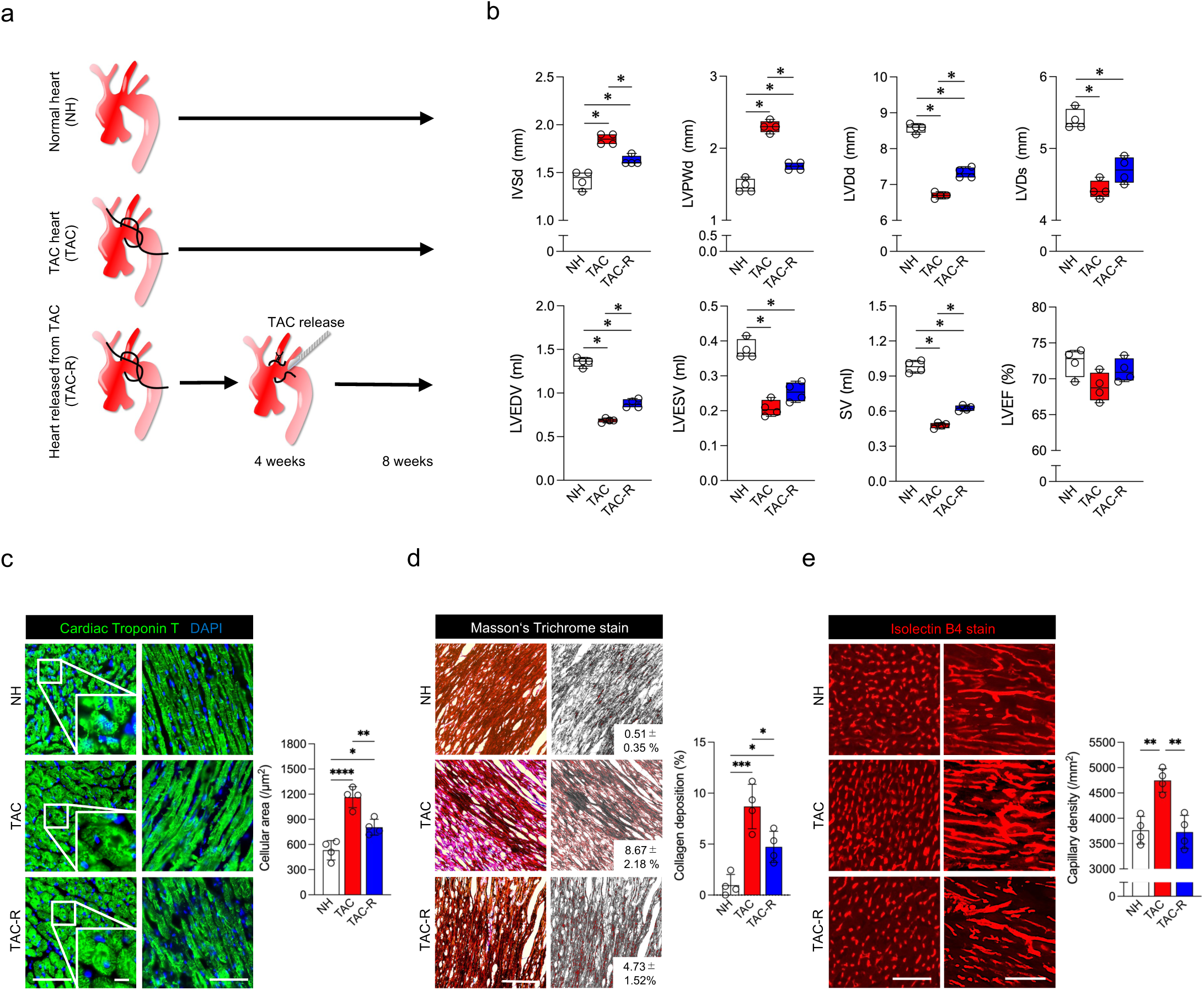
Heart released from transverse aortic constriction attenuates cardiac hypertrophy, fibrosis, and angiogenesis. (**a**) Schematic representation of the *in vivo* experiment. Transverse aortic constriction (TAC) was performed, followed by 8 weeks of observation (TAC heart). Heart was released from TAC 4 weeks after TAC surgery, reducing afterload, and this was followed by 4 weeks of observation (TAC-R). Normal heart (NH) was used for control. (**b**) Echocardiography revealed increased left ventricular posterior wall thickness at diastole (LVPWd) and decreased left ventricular end-diastolic volume (LVEDV) and left ventricular end-systolic volume (LVESV) in TAC heart relative to values in NH. Echocardiography also improvement in these changes in TAC-R heart. Other measures of interest were interventricular septal thickness at diastole (IVSd), left ventricular diastolic dimension (LVDd), left ventricular systolic dimension (LVDs), stroke volume (SV), and left ventricular ejection fraction (LVEF). n = 4 animals per group. *p < .05 versus the other group(s) by one-way ANOVA. (**c**) Cardiac troponin T staining showed a significant increase in the cross-sectional area around the short-axis of individual cardiomyocytes (viewed from the transverse section) in TAC heart. TAC release contributed to the decrease in this area in cardiomyocytes in TAC-R heart. scale bars: 50 μm in low magnification. scale bars: 10 μm in high magnification. n = 4 animals per group; mean ± SEM values are shown. *p < .05, **p < .01, ***p < .005, ****p < .001 versus the other group(s) by one-way ANOVA. (**d**) Masson’s trichrome staining showed increased collagen deposition in TAC heart. TAC release contributed to the decreased collagen deposition in TAC-R heart. (**e**) Isolectin B4 staining showed increased capillary density in TAC heart. TAC release contributed to the decreased capillary density in TAC-R heart. (**d**-**e**), scale bars: 100 μm. n = 4 animals per group; mean ± SEM values are shown. *p < .05, **p < .01, ***p < .005, ****p < .001 versus the other group(s) by one-way ANOVA.

Expression of genes associated with myocardial hypertrophy, fibrosis, and angiogenesis was examined next. Representative genes associated with cardiac hypertrophy (*Igf1, Mybpc3, Tnni3, Tnnt2, Tpm1, Actc1, Myl2,* and *Myl3*) were upregulated in TAC-subjected heart in comparison to expression of the same genes in normal heart. Expression of these genes was reduced in heart released from TAC (Fig. 2a). This trend in gene expression was similar for genes associated with fibrosis and angiogenesis. Expression of representative genes associated with fibrosis (*Col1a1, Col3a1, Mmp2, Mmp9, Tgfb1, Acta2, Timp1,* and *Fn1*) and angiogenesis (*Vegfa, Fgf1, Fgf2, Fgf9, Pdgfb,* and *Angpt2*) was upregulated in TAC-subjected heart in comparison to expression in normal heart, whereas expression of these genes was downregulated in heart released from TAC (Fig. 2b, c). Expression of genes that encode inflammatory cytokines TNFα and IL1b, which play a role in promoting transcription of myocardial matrix metalloproteinases (MMPs) and releasing latent Tgfb1 bound to the ECM, was similarly upregulated in TAC-subjected heart and downregulated in heart released from TAC (Fig. 2d). Furthermore, microarray analysis revealed the molecular signature of TAC-subjected heart and heart released from TAC against that of normal heart. Genes with a twofold or greater increase or decrease in expression between conditions were entered into Metascape, a gene annotation and analysis resource (https://metascape.org/gp/index.html#/main/step1)^14^. A representative subset of terms from the entire cluster was selected and transformed into a network layout, which characterized changes in gene expression associated with muscle hypertrophy in response to stress, muscle development, extracellular matrix organization, and blood vessel development to reflect histologic changes in TAC-subjected heart and heart released from TAC (Supplementary Fig. 1; the full dataset is available in the Gene Expression Omnibus [GEO] repository (GSE270186).

**Figure 2.**
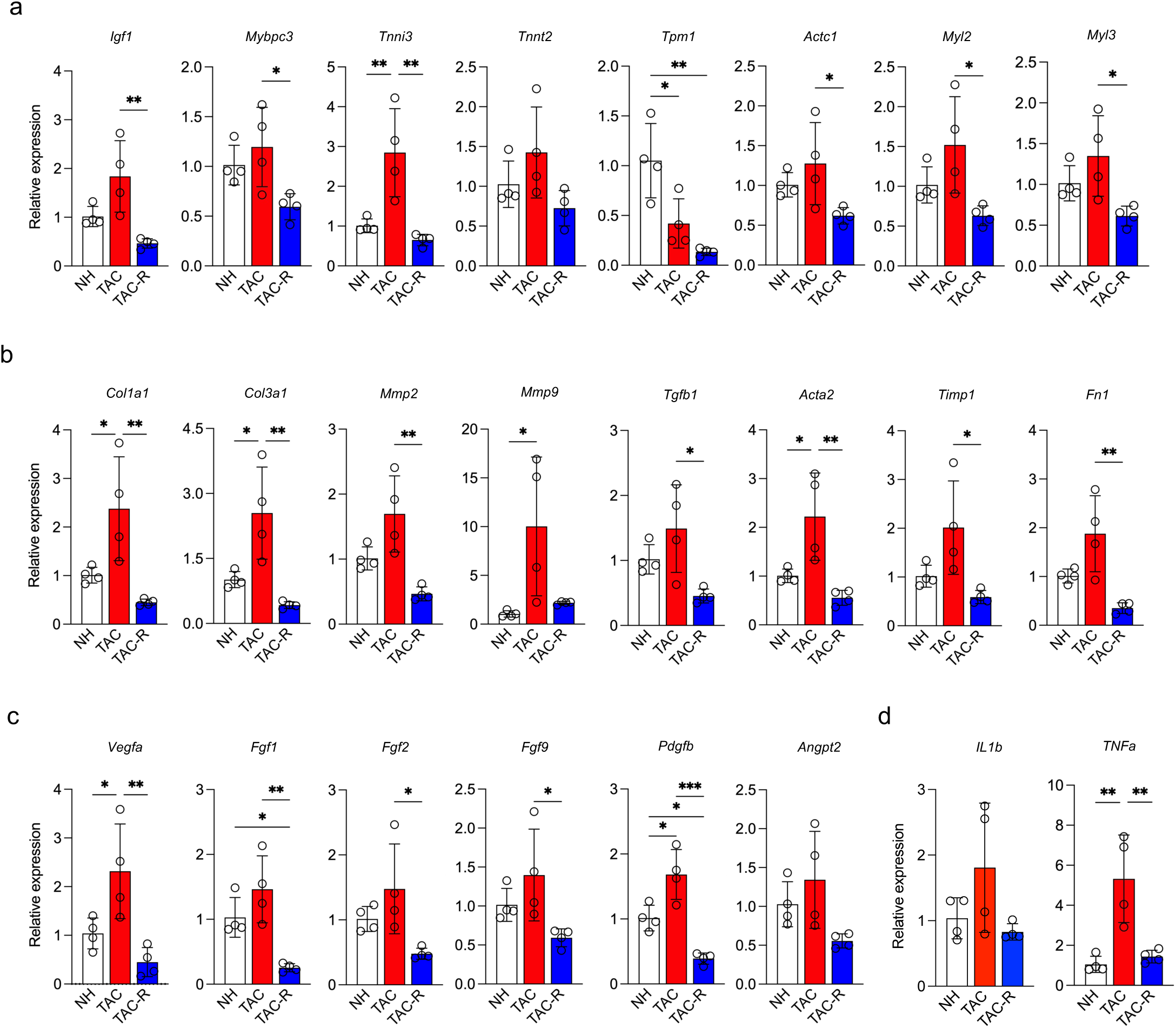
Gene expression analysis pertaining to cardiac hypertrophy, fibrosis, and angiogenesis. (**a**) qRT-PCR showed hypertrophy-associated genes (*Igf1, Mybpc3, Tnni3, Tnnt2, Actc1, Myl2,* and *Myl3*) to be upregulated, most (but not all) significantly upregulated in TAC heart relative to expression in normal heart (NH). Expression of these genes was then shown to be decreased in TAC-R heart relative to expression in NH. (**b**) qRT-PCR showed fibrosis-associated genes (*Col1a1, Col3a1, Mmp2, Mmp9, Tgfb1, Acta2, Timp1,* and *Fn1*) to be upregulated, some (but not all) significantly upregulated in TAC heart. The trend was similar in TAC-R heart. (**c**) qRT-PCR showed angiogenesis-associated genes (*Vegfa, Fgf1, Fgf2, Fgf9, Pdgfb,* and *Angpt2*) to be upregulated, some (but not all) significantly upregulated, in TAC heart. These genes were shown to be downregulated, some significantly downregulated, in TAC-R heart. (**d**) qRT-PCR showed inflammation-associated genes (*IL1b* and *TNFa*) to be upregulated, some significantly upregulated, in TAC heart. These genes were shown to be downregulated in heart TAC-R heart. (**a**-**d**), n = 4 animals per group; mean ± SEM values are shown. *p < .05, **p < .01, ***p < .005, ****p < .001 versus the other group(s) by one-way ANOVA.

### Effects of oxidative stress following reperfusion

Histologic features and expression of genes associated with myocardial hypertrophy, fibrosis, and angiogenesis were significantly altered in TAC-subjected heart relative to those in normal heart. However, these changes were reversed in heart released from TAC, with histologic features and gene expression approximating those in normal heart. Based on these results and the echocardiography-determined features, heart released from TAC appeared similar in phenotype to that of normal heart. Interestingly, principal component analysis revealed a difference in gene expression trends between normal heart and heart released from TAC, differences that were as far apart as the differences in gene expression trends between normal heart and TAC-subjected heart (Fig. 3a). Metascape was searched for relevant Gene Ontology (GO) biological process terms^14^ to determine which biological processes are involved in the changes from TAC-subjected heart to heart released from TAC, with a focus on “response to oxidative stress (GO: 0006979)” in heart released from TAC (Fig. 3b). The resolution of left ventricular afterload by TAC release appeared to improve myocardial diastolic dysfunction, increase diastolic coronary blood flow, and increase oxygen supply to the tissues. Tissue damage due to increased oxygen supply from a state of relative chronic hypoxia to re-oxygenation (reperfusion injury) was considered the main cause of elevated oxidative stress levels in heart released from TAC. Oxidative stress during such reperfusion injury is likely due in part to overproduction of reactive oxygen species (ROS). Mitochondria are the major source of ROS production. Mitochondrial ROS (mtROS) account for the majority of ROS produced in the cell. mtROS causes damage to the mitochondria themselves and increases permeability of the mitochondrial membrane. This causes a rapid influx of calcium and other molecules into and out of the mitochondria, inducing mitochondrial dysfunction^15–17^. mtDNA copy number, a measure of mitochondrial function, was therefore assessed and found to be reduced in heart released from TAC more than in TAC-subjected heart, suggesting association between the increase in oxidative stress following reperfusion injury and decreased mitochondrial function (Fig. 3c). Heatmaps generated by a group of genes associated with the GO biological process term “response to oxidative stress” revealed markedly accelerated expression of regulator of calcineurin 1 (*Rcan1*), especially in heart released from TAC (Fig. 3d), a finding confirmed by quantitative reverse transcription-polymerase chain reaction (qRT-PCR) (Fig. 3e). RCAN1 is primarily responsible for (i) maintenance of mitochondrial function^18^, (ii) inhibition of inflammation^19^, (iii) inhibition of Rho kinase activity^20^, and (iv) inhibition of calcineurin-NFAT signaling^20^. In this context, enrichment analysis was performed to clarify the role of *Rcan1* in the oxidative stress state associated with reperfusion (the TAC-release condition). Genes with a twofold or greater increase in expression in heart released from TAC compared to expression in TAC-subjected heart were entered into Enricher, a comprehensive resource for curated gene sets as well as a search engine (https://maayanlab.cloud/Enrichr/)^21^. The GO biological process terms related to (i)–(iv) noted above were identified (Table 1). Of these, negative regulation of calcineurin-NFAT signaling cascade (GO:0070885) was found to be a biological process closely (though not significantly) related to the change from the TAC condition to the TAC-release condition. The calcineurin-NFAT signaling cascade is a stress response in which calcineurin activates NFAT transcription factors, leading to the expression of genes responsible for hypertrophy, fibrosis, and angiogenesis. Activation of calcineurin-NFAT signaling by oxidative stress simultaneously induces its own negative regulator, RCAN1^19^. The data obtained indicate that activation of the calcineurin-NFAT signaling cascade due to oxidative stress induced by reperfusion following absence of afterload in heart released from TAC is inhibited by RCAN1. In other words, findings suggest the existence of a cardiac-specific homeostatic mechanism effecting hypertrophy, fibrosis, and angiogenesis, with RCAN1 as one of the key regulators (Fig. 3f).

**Figure 3.**
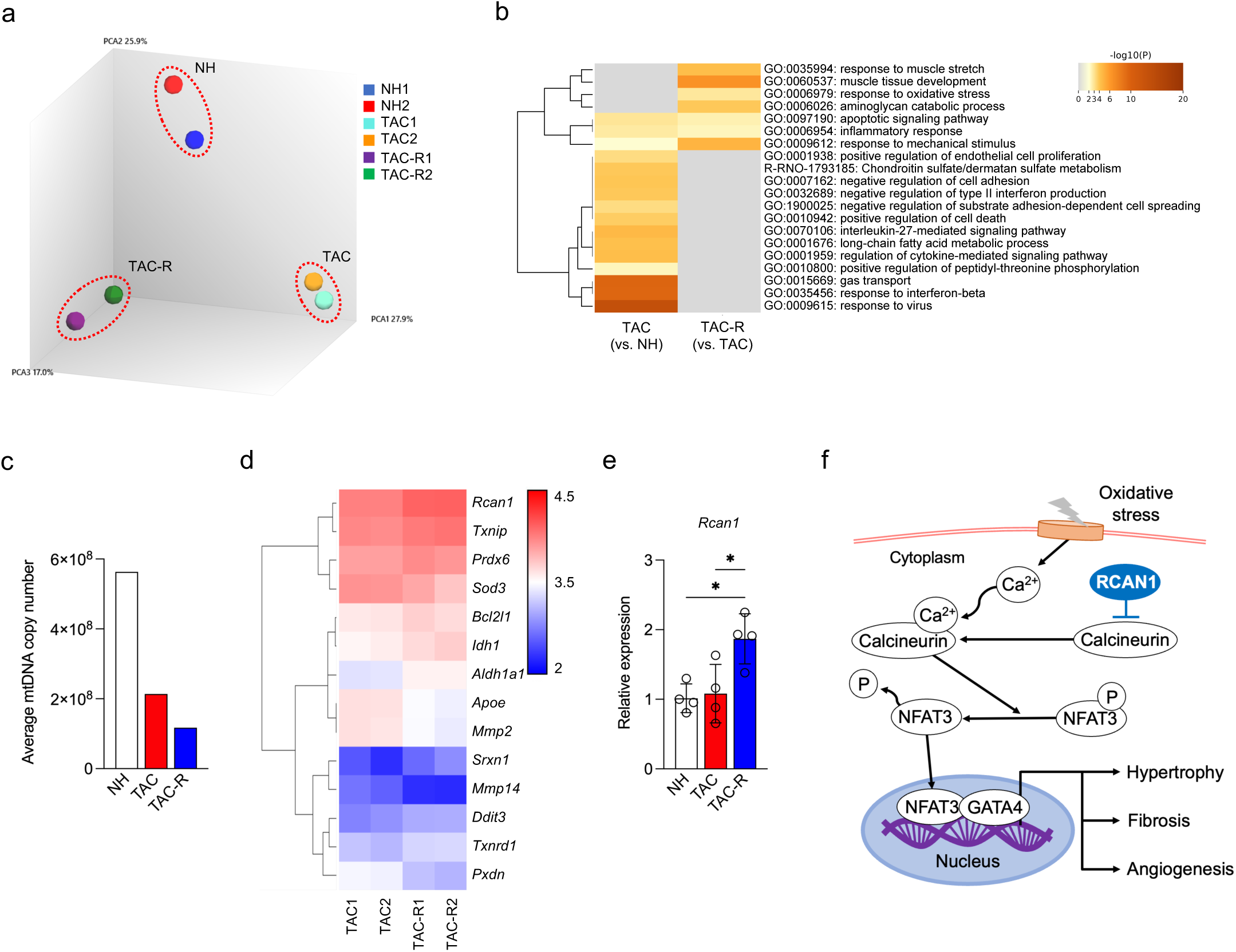
Reperfusion following left ventricular loading increases oxidative stress. (**a**) Principal component analysis clearly revealed distinct grouping of normal heart (NH), TAC heart, and TAC-R heart, indicating that TAC and TAC-R hearts have unique gene expression patterns compared with NH. (**b**) The top 20 significantly enriched molecules are displayed in the heatmap. The heatmap cells are colored according to their p-values, with white indicating lack of enrichment for that term in the corresponding gene list. (**c**) Average mitochondrial DNA (mtDNA) copy number showed mitochondrial function to be significantly attenuated in TAC heart. In TAC-R heart, average mtDNA copy number was further decreased. (**d**) Comprehensive gene expression analysis showed significant changes in the expression of 14 genes associated with Gene Ontology term “Response to oxidative stress” including *Rcan1*. (**e**) qRT-PCR showed revealed *Rcan1* to be significantly upregulated in TAC-R heart. n = 4 animals per group; mean ± SEM values are shown. *p < .05 versus the other group(s) by one-way ANOVA. (**f**) Schematic representation and overview of the calcineurin-nuclear factor of activated T cells (NFAT) signaling pathway. NFAT proteins are typically located in the cytoplasm in a phosphorylated, inactive state. Various stressors can directly activate calcineurin, prompting the dephosphorylation and subsequent nuclear translocation of NFAT. Once dephosphorylated by calcineurin, NFAT proteins move from the cytoplasm to the nucleus, where they act as transcription factors regulating genes involved in myocardial hypertrophy, fibrosis, and angiogenesis.

**Table 1.** Gene Ontology (GO) terms associated with maintenance of mitochondrial function, inhibition of inflammation, inhibition of Rho kinase activity, and inhibition of calcineurin-NFAT signaling. *Negative regulation of calcineurin-NFAT signaling cascade (GO:0070885) is more involved in the effects of release from TAC than in other biological processes.

| Name | p-value | Odds Ratio |
| --- | --- | --- |
| Regulation of mitochondrion organization (GO: 0010821) | 0.3672 | 2.23 |
| Regulation of cytokine production involved in inflammatory response (GO:1900015) | 0.3436 | 2.43 |
| Inflammatory response (GO:0006954) | 0.3636 | 1.41 |
| Positive regulation of inflammatory response (GO:0050729) | 0.6145 | 1.06 |
| Regulation of inflammatory response (GO:0050727) | 0.6446 | 0.91 |
| Rho protein signal transduction (GO:0007266) | 0.3787 | 2.14 |
| Negative regulation of calcineurin-NFAT signaling cascade (GO:0070885) | 0.0957 | 10.94 |
| Regulation of calcineurin-NFAT signaling cascade (GO:0070884)* | 0.2673 | 3.31 |

### RCA1 is a potential regulator of cardiac remodeling

To affirm the importance of *Rcan1* in the regulatory mechanism of myocardial hypertrophy, fibrosis, and angiogenesis in myocardial environments with different hemodynamic loads, hearts released from TAC were transfected *in vivo* with negative control small interfering RNA (NC siRNA) or *Rcan1* siRNA (Fig. 4a). Results were as follows: Expression of *Rcan1* was confirmed to be decreased in *Rcan1* siRNA-transfected heart (Fig. 4b). M-mode echocardiography performed 4 weeks after TAC surgery revealed a significant increase in left ventricular wall thickness and a reduction in left ventricular volume and LVEF in TAC-subjected heart compared to measures in normal heart. Subsequently, in NC siRNA-transfected heart, echocardiography performed at 8 weeks showed improvement in these alterations. However, an increase in left ventricular wall thickness and a reduction in left ventricular volume and LVEF were seen in *Rcan1* siRNA-transfected heart, compared to measures in NC siRNA-transfected heart (Fig. 4c, d). In *Rcan1* siRNA-transfected heart, the weight change ratio was significantly reduced compared to that of normal heart and of NC siRNA-transfected heart (Fig. 5a), suggesting cardiac cachexia due to reduced cardiac function as a cause. Histology was then investigated. Extracted heart weight corrected for body weight did not differ between normal heart and NC siRNA-transfected heart but was significantly increased in *Rcan1* siRNA-transfected heart (Fig. 5b). Interventricular septum thickness in NC siRNA-transfected heart was similar to that in normal heart but significantly increased in the *Rcan1* siRNA-transfected heart compared to that in both normal heart and NC siRNA-transfected heart (Fig. 5c). The trend was the same as that of left ventricular wall thickness. The cross-sectional area around the short-axis of individual cardiomyocytes (Fig. 5d), amount of collagen deposition (Fig. 5e), and capillary density (Fig. 5f) were significantly increased in *Rcan1* siRNA-transfected heart. Increased (though not always significantly increased) expression of hypertrophy-related genes (*Igf1, Mybpc3, Tnni3, Tnnt2, Tpm1, Actc1,* and *Myl3*) (Fig. 6a), fibrosis-related genes (*Col1a1, Col3a1, Mmp2, Tgfb1, Timp1,* and *Fn1*) (Fig. 6b), and angiogenesis-related genes (*Fgf1, Fgf2,* and *Angpt2*) (Fig. 6c) in heart released from TAC transfected with *Rcan1* reflected an increased cross-sectional area around the short-axis of individual cardiomyocytes, increased amount of collagen deposition, and increased capillary density. Briefly, *Rcan1* knockdown attenuated cardiac reversibility and plasticity in heart released from TAC. Thus, *Rcan1* appears to be a principal gene regulating the reversibility and plasticity of cardiomyocyte hypertrophy, fibrosis, and angiogenesis in heart released from TAC.

**Figure 4.**
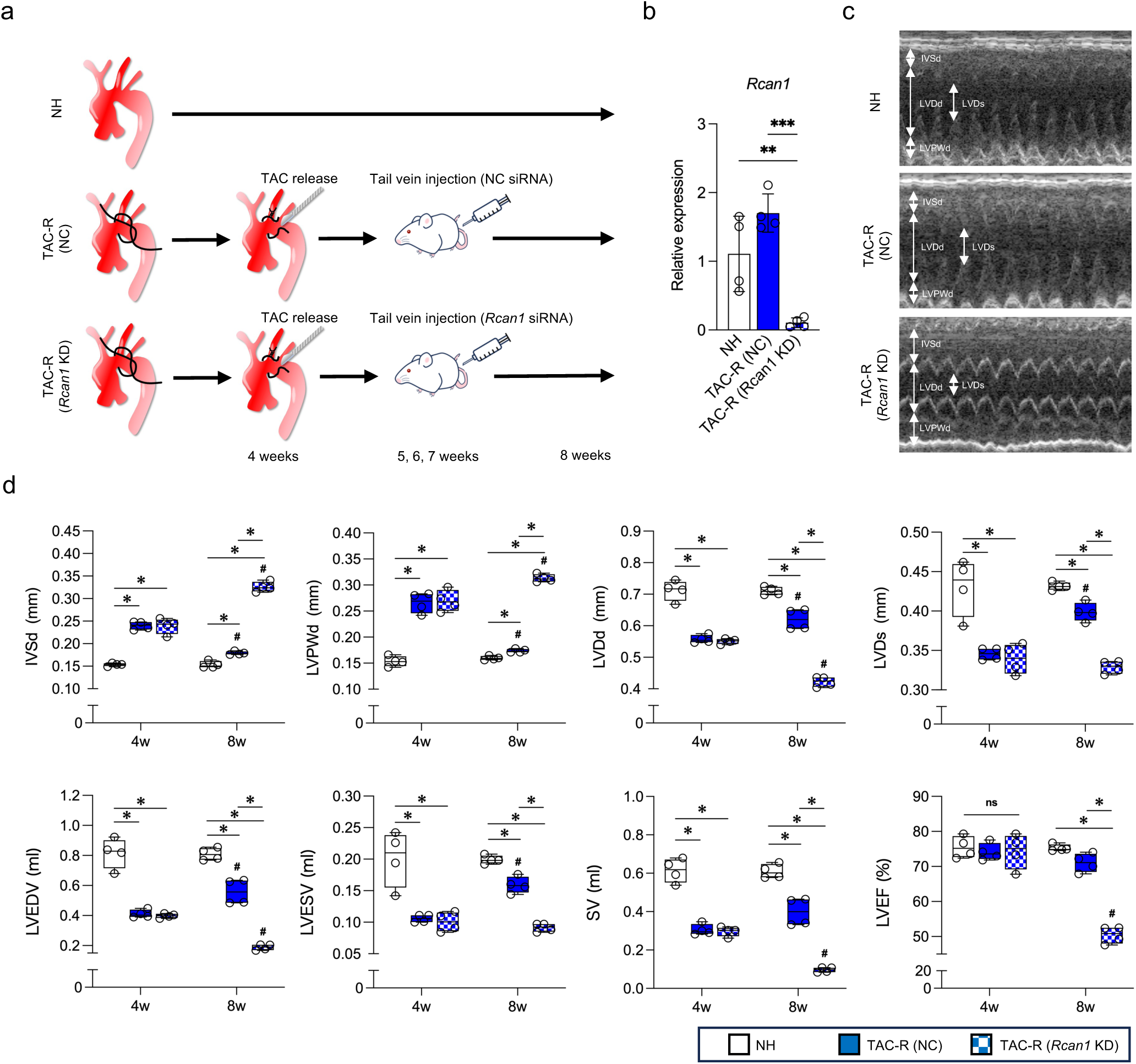
Knockdown of regulator of calcineurin 1 exacerbates hypertrophy and attenuates cardiac contractility. (**a**) Schematic representation of the *in vivo* experiment. Regulator of calcineurin 1 (*Rcan1*) siRNA or negative control (NC) siRNA combined with JetPEI^®^ was injected into the tail vein of rats 1 week after transverse aortic constriction (TAC) release (TAC-R heart), followed by a total of three intravenous infusions (one given every other week). (**b**) qRT-PCR showed *Rcan1* expression to be increased in heart released from TAC transfected with NC siRNA. In contrast, its expression was markedly suppressed in TAC-R heart transfected with *Rcan1* siRNA. n = 4 animals per group; mean ± SEM values are shown. *p < .05, **p < .01, ***p < .005, ****p < .001 versus the other group(s) by one-way ANOVA. (**c**) Representative M-mode echocardiography tracings of normal heart (NH) and TAC-R heart transfected with NC *Rcan1* siRNA or *Rcan1* siRNA. (**d**) Echocardiography revealed increased left ventricular wall thickness and decreased left ventricular end diastolic and end systolic volumes (LVEDV and LVESV) in TAC heart at 4 weeks after TAC surgery, and improvement in such changes in TAC-R heart transfected with NC siRNA at 8 weeks after surgery. *Rcan1* knockdown further increased left ventricular posterior wall thickness and decreased left ventricular volume, and it attenuated LVEF in TAC-R heart examined at 8 weeks after surgery. n = 4 animals per group. *p < .05 versus the other group(s) by one-way ANOVA; ^#^*P*<0.05 versus the same group at 4 weeks after surgery by paired t-test. TAC-R (NC), heart released from TAC transfected with NC siRNA; TAC-R (*Rcan1* KD), heart released from TAC transfected with *Rcan1* siRNA; IVSd, interventricular septal thickness at diastole; LVPWd, left ventricular posterior wall thickness at diastole; LVDd, left ventricular diastolic dimension; LVDs, left ventricular systolic dimension; LVEDV, left ventricular end-diastolic volume; LVESV, left ventricular end-systolic volume; SV, stroke volume; EF, ejection fraction.

**Figure 5.**
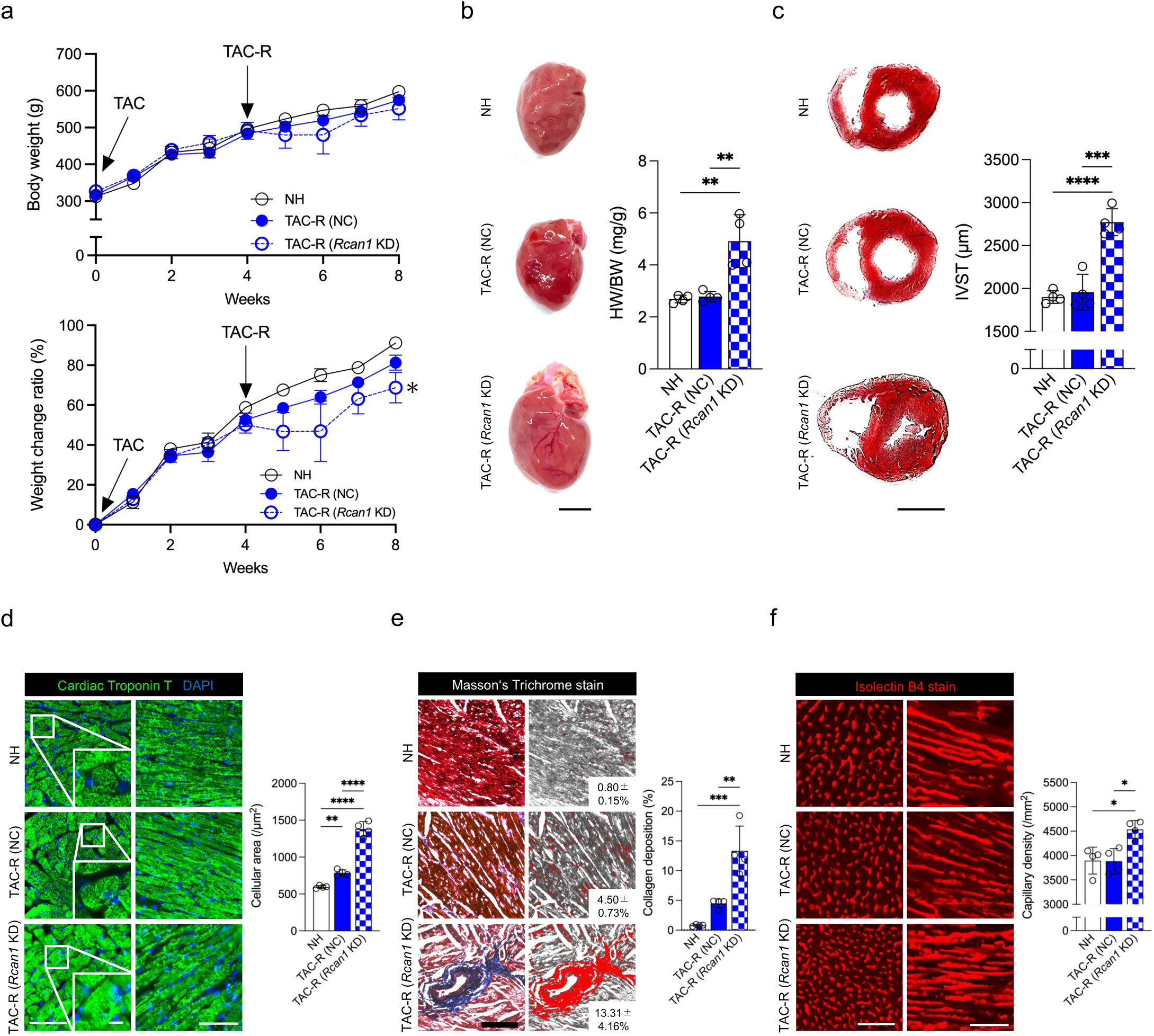
Regulator of calcineurin 1 knockdown (KD) exacerbates hypertrophy, fibrosis and angiogenesis in heart. (**a**) Body weight change (upper graph) and its ratio (lower graph) of normal heart (NH), Heart released from TAC transfected with negative control (NC) siRNA (TAC-R [NC]) or *Rcan1* siRNA (TAC-R [*Rcan1* KD]). (**b**) Representative images of NH and TAC-R (NC) and TAC-R (*Rcan1* KD) hearts. Heart weight (HW)/body weight (BW) ratio (mg/g) was markedly increased in TAC-R (*Rcan1* KD) heart. (**c**) Representative images of the cross-sectional area around the short-axis of individual cardiomyocytes (viewed from the transverse section) of NH and TAC-R (NC), and TAC-R (*Rcan1* KD) hearts. Interventricular septal thickness (IVST) was markedly increased in TAC-R (*Rcan1* KD) heart. (**d**) Cardiac troponin T staining showed significantly increased area of short-axis transverse section of cardiomyocyte in TAC-R (*Rcan1* KD) heart. (**e**) Masson’s trichrome staining showed significantly increased collagen deposition in TAC-R (*Rcan1* KD) heart. (**f**) Isolectin B4 staining showed increased capillary density in TAC-R (*Rcan1* KD) heart. (a) n = 4 animals per group. *p < .05 versus the other group(s) by one-way ANOVA; (b, c) scale bars: 10 mm; (d) scale bars: 50 μm in low magnification. scale bars: 10 μm in high magnification; (e, f) scale bars: 100 μm; (**a–e**) n = 4 animals per group; mean ± SEM values are shown. *p < .05, **p < .01, ***p < .005, ****p < .001 versus the other group(s) by one-way ANOVA.

**Figure 6.**
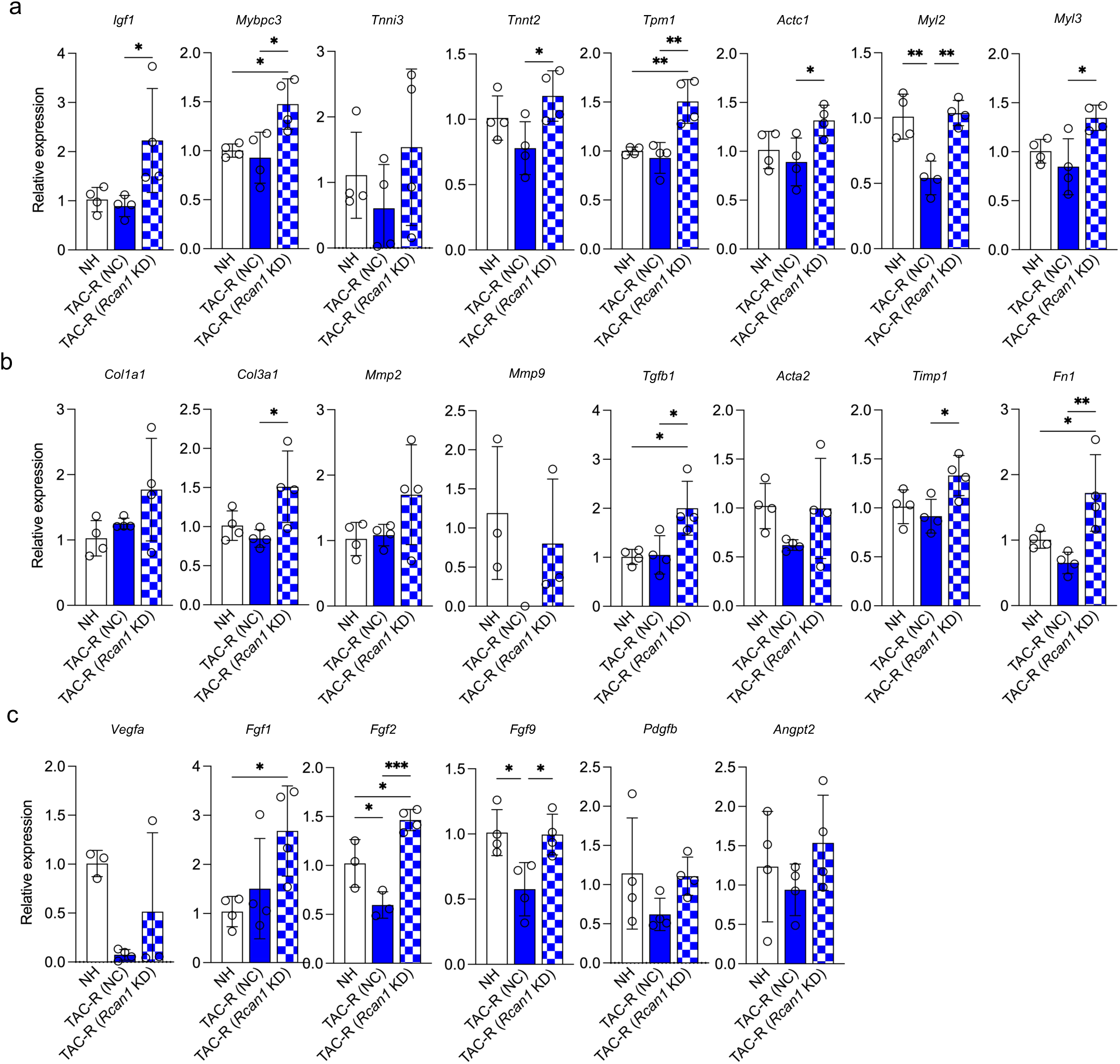
Knockdown of regulator of calcineurin 1 affects expression of genes related to cardiac hypertrophy, fibrosis, and angiogenesis. (**a**) qRT-PCR revealed upregulated expression of hypertrophy-associated genes (*Igf1, Mybpc3, Tnnt2, Tpm1, Actc1, Myl2* and *Myl3*) in TAC-R heart transfected with *Rcan1* siRNA (TAC-R [*Rcan1* KD]) relative to expression in normal heart and TAC-R heart transfected with negative control (NC) siRNA (TAC-R [NC]). (**b**) qRT-PCR revealed significantly upregulated expression of fibrosis-associated genes (*Col3a1, Tgfb1, Timp1,* and *Fn1*) in TAC-R (*Rcan1* KD) heart relative to that in normal heart and/or TAC-R (NC) heart. (**c**) qRT-PCR revealed significantly upregulated expression of angiogenesis-associated genes (*Fgf1, Fgf2,* and *Fgf9*) in TAC-R (*Rcan1* KD) heart relative to that in normal heart and/or TAC-R NC heart. (**a–c**) n = 4 animals per group; mean ± SEM values are shown. *p < .05, **p < .01, ***p < .005, ****p < .001 versus the other group(s) by one-way ANOVA.

## Discussion

Cardiac hypertrophy frequently causes structural and functional modifications in the heart, including changes in coronary blood flow, which can lead to reduced flow due to diastolic failure and impaired myocardial perfusion^22^. Resolution of the hypertrophy, whether via therapeutic interventions or adaptive mechanisms, can lead to reperfusion of previously ischemic or hypoxic myocardial tissue. Although reperfusion restores blood flow to these areas, it can also induce oxidative stress and tissue damage through various mechanisms, such as ROS overproduction and inflammation^23^. Inflammatory cells and mediators contribute to oxidative stress through pathways including nicotinamide adenine dinucleotide phosphate oxidase activation and ROS release^24^. Additionally, cardiac hypertrophy is linked to changes in mitochondrial function^25^. Resolving hypertrophy may enhance mitochondrial function, but it can temporarily disrupt mitochondrial homeostasis, leading to ROS production and oxidative stress^26^. In myocardium, ROS are constantly produced by a variety of enzymes, including mitochondrial electron transfer enzymes, and under physiological conditions ROS in myocardial tissue are maintained at low levels by intrinsic antioxidant mechanisms. However, during reperfusion injury, overproduction of ROS results in a state of oxidative stress^15^. The ROS induce opening of the mitochondrial permeability transition pore, which leads to cell death by activation of the apoptotic pathway, resulting in mitochondrial dysfunction, which in turn produces mtROS^17^. Indeed, mtROS production during reperfusion injury correlates with reduced myocardial function and myocyte death^16^. The study described herein revealed the following as a possible mechanism inducing oxidative stress under TAC release: (1) reversal of the physical stenosis created by TAC decreasing left ventricular afterload and increasing coronary reperfusion; (2) increased oxidative stress through the production of ROS; and (3) mitochondrial dysfunction resulting in the production of mtROS, further contributing to the oxidative stress.

The calcineurin-NFAT (nuclear factor of activated T cells) signaling pathway that was suggested by results of the enrichment analysis to relate to phenotypic change is integral to various physiological processes, including immune response and oxidative stress. Calcineurin, a calcium/calmodulin-dependent phosphatase, activates NFAT transcription factors, leading to the expression of genes responsible for cardio-structural alterations. NFAT proteins are typically located in the cytoplasm in a phosphorylated, inactive state. ROS can directly activate calcineurin, prompting dephosphorylation and subsequent nuclear translocation of NFAT proteins. Once dephosphorylated by calcineurin, NFAT proteins move from the cytoplasm to the nucleus, where they act as transcription factors regulating genes involved in myocardial hypertrophy^27^. Activation of the calcineurin-NFAT pathway by ROS or oxidative stress contributes to pathological cardiac remodeling and heart failure.

Genes such as *Igf1* (insulin-like growth factor 1)^28^, *Mybpc3* (myosin binding protein C)^29^, *Tnni3* (troponin I)^30^, *Tnnt2* (troponin T)^31^, *Tpm1* (tropomyosin 1)^31^, *Actc1* (actin alpha cardiac muscle 1)^32^, *Myl3* (myosin light chain 3)^33^, and *Myl2* (myosin light chain 2)^34^ are crucial for cardiac hypertrophy. In response to various stimuli, expression of these genes is upregulated, causing cardiac hypertrophy and remodeling. In a comparison of normal heart, TAC-subjected heart, and heart released from TAC, significant up-regulation of *Tnni3* was observed in TAC-subjected heart and related to cardiac hypertrophy. Up-regulation, though not significant, of *Igf1, Mybpc3, Tnnt2, Actc1, Myl2,* and *Myl3* was also observed. In comparison to both normal heart and NC siRNA-transfected heart, *Rcan1* siRNA-transfected heart showed significantly increased expression of *Mybpc3*, related to increased cardiac hypertrophy. Expression of *Igf1, Tnnt2, Tpm1, Actc1,* and *Myl3* tended to be increased. The calcineurin-NFAT signaling pathway is crucial for transcriptional regulation of *Igf1*^35^ and *Mybpc3*^36^. However, transcriptional regulation of other cardiac hypertrophy-related genes by the calcineurin-NFAT signaling pathway was not clarified. The author speculates that *Rcan1* knockdown in heart released from TAC affected the expression of other cardiac hypertrophy-related genes, including *Igf1* and *Mybpc3*, whose transcriptional regulation mechanisms are already known, mainly by enhancing the calcineurin-NFAT signaling pathway. To prove this, studies are needed to clarify the association between transcriptional regulation of cardiac hypertrophy-related genes other than *Igf1* and *Mybpc3* and the calcineurin-NFAT signaling pathway.

With the ECM being an essential component of myocardium, alterations in the ECM significantly affect myocardial structure and function. The calcineurin-NFAT signaling pathway regulates genes involved in cardiac fibrosis, including collagen genes such as *Col1a1* (collagen type I alpha 1 chain), *Col3a1* (collagen type III alpha 1 chain), *Mmp2* (matrix metallopeptidase 2), and *Mmp9* (matrix metallopeptidase 9)^37, 38^. This pathway also influences *Tgfb1* (transforming growth factor beta 1), *Acta2* (actin alpha 2, smooth muscle)^37^. Furthermore, inflammatory cytokines such as TNFa (tumor necrosis factor alpha) and IL1b (interleukin-1 beta) drive the transcription of MMPs in the myocardium^39^, releasing latent TGFB1 bound to the ECM. Therefore, besides the calcineurin-NFAT signaling pathway, increased expression of inflammatory cytokines in TAC-subjected heart and its decreased expression in heart released from TAC may have affected the regulation of MMP and TGFB1 expression. TGFB1 then activates the Smad signaling pathway in fibroblasts, promoting the production of fibrous collagens type I and III, key components of the ECM^40–42^. ACTA2, a marker of activated myofibroblasts, is crucial in cardiac fibrosis, enhancing myofibroblast differentiation and the contractile and matrix-producing abilities of fibroblasts. FN1 (fibronectin 1), another significant ECM protein, is upregulated in cardiac fibrosis. Because MMP2 and MMP9 are downstream factors of ROS and inflammatory cytokines TNFa and IL-1b, their expression and activity increase during myocardial remodeling, promoting interstitial fibrosis. Studies on MMP2-deficient mice have shown reduced myocardial hypertrophy and fibrosis in response to hypertension and cardiac pressure overload^43^. Similarly, MMP9 deficiency partially improves myocardial hypertrophy and fibrosis after pressure overload^44^, pointing to the pivotal roles of MMP2 and MMP9 in fibrosis and ECM remodeling. MMPs are regulated by tissue metalloproteinase inhibitors (TIMPs), and ECM remodeling is intricately related to the MMP/TIMP balance^45^. Left ventricular fibrosis may result from both increased collagen synthesis and reduced collagen degradation due to decreased collagenase activity, which is associated with increased *Timp1* expression^46^. In TAC-subjected heart, increased collagen synthesis and changes in *Timp1* and *Mmp* expression and the MMP/TIMP balance may have promoted collagen deposition. Conversely, in heart released from TAC, changes in expression of *Mmps* and *Timp1* and the balance between them may have suppressed left ventricular fibrosis. MMPs and TIMPs may also individually influence fibroblast behavior and ECM production. Therefore, other regulatory mechanisms governing diverse fiber production need to be elucidated for us to understand myocardial fibrosis.

The calcineurin-NFAT signaling pathway governs expression of essential angiogenic factors, including VEGFA (vascular endothelial growth factor A)^47^, FGF1 (fibroblast growth factor 1)^48^, PDGFB (platelet-derived growth factor beta)^49^, and ANGPT2 (angiopoietin 2)^50^. VEGFA is a potent inducer of angiogenesis, the process by which new blood vessels form from existing ones. In cardiac hypertrophy, increased *Vegfa* expression leads to increased levels of VEGFA, stimulating growth of new blood vessels in the heart. Whereas such angiogenesis initially meets the heightened oxygen and nutrient demands of hypertrophic cardiac tissue, prolonged angiogenesis can contribute to adverse remodeling and heart failure^51^. Thus, regulation of VEGFA by the calcineurin-NFAT pathway plays a crucial role in balancing beneficial and detrimental angiogenesis occurring with cardiac hypertrophy. FGF1, FGF2, and FGF9 are key angiogenic factors that bind to fibroblast growth factor receptors, activating intracellular signaling cascades^52^. Such binding promotes blood vessel formation and endothelial cell proliferation, ensuring adequate blood supply to the expanding myocardium. PDGFB is crucial for pericyte recruitment and vascular stabilization during angiogenesis. Secreted by endothelial cells and other myocardial cells, PDGFB binds to receptors on pericytes and vascular smooth muscle cells, promoting their recruitment to newly formed blood vessels^53^. ANGPT2 is a context-dependent angiogenic factor that modulates vessel remodeling and maturation. Acting as a competitive antagonist to ANGPT1, ANGPT2 disrupts the TIE2 receptor signaling pathway and destabilizes endothelial cell junctions. This destabilization promotes vessel sprouting and remodeling, facilitating angiogenesis in the myocardium^54^. Collectively, VEGFA, FGF, PDGFB, and ANGPT2 play critical roles in regulating angiogenesis, ensuring proper vascular adaptation with development of cardiac hypertrophy. The study reported herein revealed a certain link between the transcriptional regulation of angiogenesis-related genes and the calcineurin-NFAT signaling pathway in response to hemodynamic changes, but further studies are needed.

*Rcan1* gene, known as regulator of calcineurin 1 gene, is a crucial inhibitor of the calcineurin-NFAT signaling pathway, and it plays a role in preventing cardiac hypertrophy^20^. Overexpression of RCAN1 has been shown to attenuate pathological cardiac injury in ischemia/reperfusion animal models by suppressing calcineurin-NFAT signaling and hypertrophic gene expression. Conversely, decreased expression of RCAN1 is associated with exacerbated cardiac injury^55^. Studies have shown that *Rcan1* expression is dynamically regulated by oxidative stress during inflammation^19^. The present study on oxidative stress through increased coronary blood flow with reduced left ventricular afterload and the regulation of calcineurin-NFAT signaling activity by RCAN1 provided valuable insight into the molecular mechanisms underlying cardiac reversibility and plasticity. RCAN1 can function as a pro-survival factor, protecting cells from stress-induced apoptosis by inhibiting calcineurin-NFAT signaling activation^18^. Thus, during increased oxidative stress associated with reperfusion, RCAN1 may determine the balance between cell survival and cell death, not just effect cardiac tissue remodeling. Optimal regulation of RCAN1 expression and activity is essential for coordinating the cellular responses necessary for reversibility and plasticity mechanisms in cardiac hypertrophy.

Understanding the role of ROS generation and oxidative stress due to reperfusion injury following relief from left ventricular afterload has important therapeutic implications for cardiac repair and regeneration strategies. Targeting oxidative stress or modulating RCAN1 expression/activity through gene therapy or pharmacological interventions could be potential therapeutic approaches to attenuate cardiac hypertrophy and fibrosis and thereby improve cardiac function. Further research in the form of preclinical and clinical studies is needed to elucidate specific mechanisms and explore the therapeutic potential of targeting this axis.

## Limitations

In the study described, markers of cardiac hypertrophy, fibrosis, and angiogenesis were assessed by qRT-PCR, with total RNA from whole hearts used as starting material. Therefore, the results do not allow for a conclusion whether the transcriptional differences originate from cardiomyocytes, fibroblasts, or endothelial cells. Further studies are needed to isolate each cell type from normal heart, TAC-subjected heart, and heart released from TAC and then run a cell-by-cell analysis to determine the role of each cardiac component cell. Promotion of the calcineurin-NFAT signaling pathway by suppression of RCAN1 expression was found to have a significant effect on the cardiac tissue recovery process, i.e., suppression of RCAN1 expression may exacerbate cardiac hypertrophy and fibrosis and promote angiogenesis. However, besides regulating the calcineurin-NFAT signaling pathway, RCAN1 also regulates other pathways involved in cardiac hypertrophy, fibrosis, and angiogenesis, including the RHO and TGFb (transforming growth factor-beta) signaling pathways. Thus, it is uncertain whether exacerbation of cardiac hypertrophy and fibrosis by *Rcan*1 knockdown in the TAC-release model results solely from enhanced calcineurin-NFAT signaling.

## Conclusion

A left ventricular afterload relief model was established by releasing TAC, and study of the model shed light on the structural, functional, and histologic changes that occur in the heart during the recovery from pressure overload-induced cardiac hypertrophy (Fig. 7). Increased oxidative stress upon reperfusion and upregulation of the *Rcan1* gene, suppressing the calcineurin-NFAT signaling pathway, are pivotal in the tissue repair process following heart damage. Their intricate interaction dynamically influences cellular responses to injury, inflammation resolution, and tissue remodeling. These study findings provide insight into cardiac repair and regeneration. Further exploration in this realm may ultimately open novel therapeutic avenues for managing heart disease.

**Figure 7.**
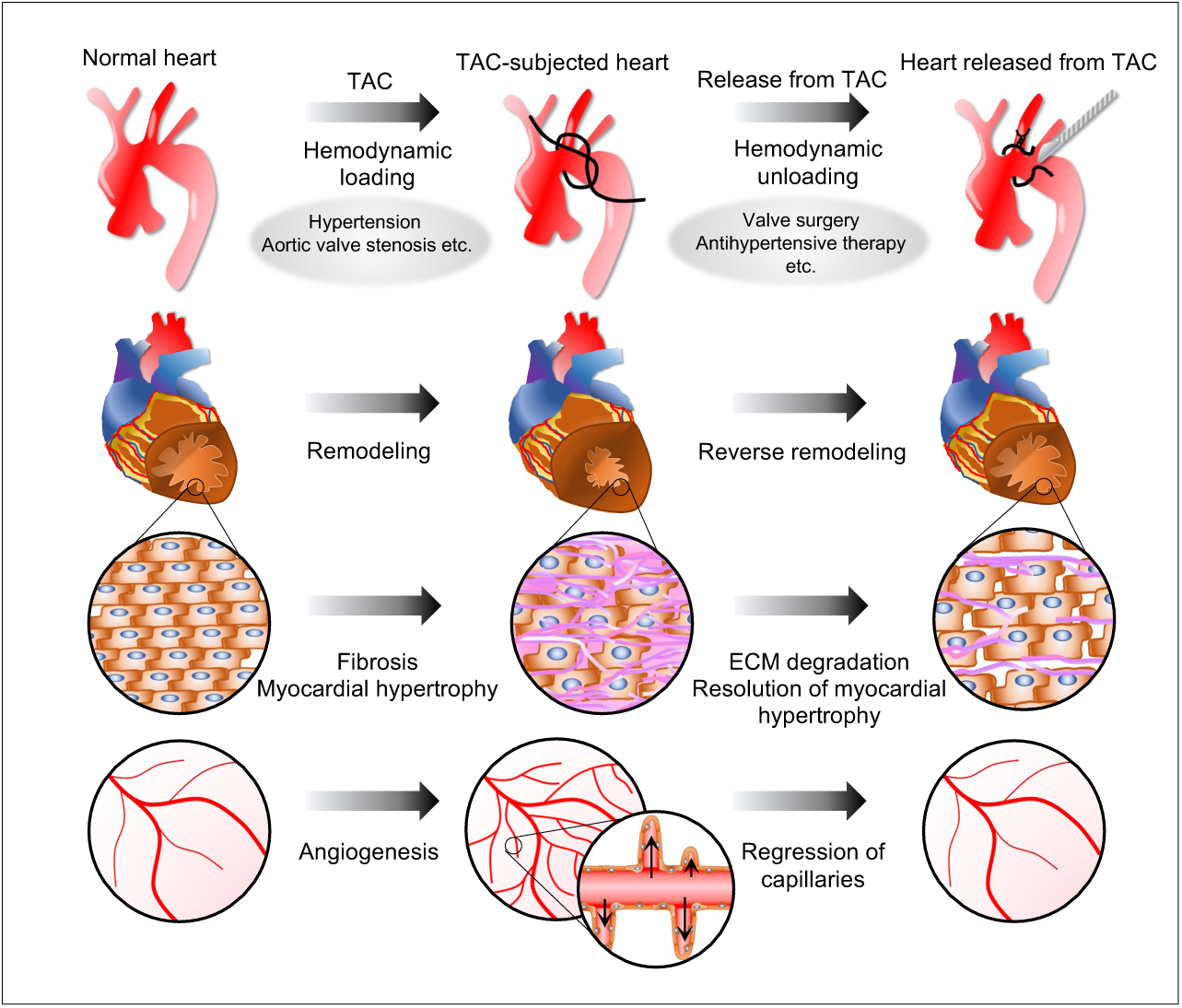
Schematic representation and overview of reversibility of hemodynamic changes in the heart. Increased left ventricular afterload resulting from transverse aortic constriction (TAC) promotes cardiac hypertrophy, fibrosis, and angiogenesis. In contrast, reduction in afterload by TAC release (TAC-R) attenuates these changes. The structural change in the heart involves modulation of the activity of the calcineurin nuclear factor of activated T cells (NFAT) signaling pathway, with a potential regulator of calcineurin 1.

## Materials and methods

### TAC procedure and TAC release

Animal experiments were conducted on approval of the Animal Experiment Committee in accordance with the Animal Experiment Regulations of the Center for Experimental Medicine, Jichi Medical University (Approval No. 21058-01). TAC surgery and TAC-release were performed as previously reported but with some modifications^13, 56–58^. Animals used were male Sprague Dawley rats (10 weeks old). Food and water were withheld 6 hours prior to surgery. Surgical instruments were sterilized in an autoclave, and the surgical field was disinfected with 75% isopropyl alcohol. A heating pad was used during surgery, set to an appropriate temperature (37℃) to maintain the animals’ normal body temperature and thereby avoid rapid changes in heart and respiratory rates. Inhalation anesthesia was achieved with a mixture of 2% isoflurane and 100% oxygen at 0.5–1.0 L/min. Hairs on the animal’s anterior neck and chest were removed with depilatory cream. Each rat was positioned supine on the heating pad, with their anterior teeth fixed over silk threads, the neck stretched, and limbs secured with paper surgical tape. Anesthesia was maintained throughout the surgery by administration of 1.5–2% isoflurane and 0.5–1.0 L/min 100% oxygen and confirmed by the toe-pinch test. The surgical field was sterilized with 70% alcohol, and sterile drapes and gloves were used for each rat. Skin incisions were made with a scalpel at the midline of the rat’s neck and chest. The muscular layer over the trachea was separated bilaterally. Partial sternotomy, up to the second rib, was performed, and the sternum was separated with a chest retractor. The thymus was removed and adipose tissue was gently detached from the aortic arch. After identification of the transverse aorta, a 3-0 silk thread was passed between the brachiocephalic artery and left common carotid artery with the use of sharp forceps. A loop of about 10 mm in diameter was created with 5-0 monofilament thread, and a 3-0 silk thread was passed through it (to indicate where to later cut the silk thread for TAC-release). A 21-gauge blunt needle was placed parallel to the aorta, the 3-0 silk thread was quickly tied to prevent loosening, and the needle was removed to produce aortic stenosis. In control rats, procedures were the same, with the exception of aortic ligation. The chest traction device was removed, the sternum was brought together with 4-0 silk thread, and the skin was closed by continuous suturing with 6-0 monofilament. Postoperative analgesia was provided by means of intraperitoneal buprenorphine injection. Anesthesia was gradually lessened, and the rats were allowed to recover on a heating pad on signs of spontaneous respiration. After full recovery, the rats were returned to their original housing room under a 12-hour light/dark cycle and fed soft food. For pain relief, rats were given buprenorphine (0.1 mg/kg) intraperitoneally every 12 hours for 3 days. Four weeks after the transverse aortic ligation, skin incisions were made at the neck and chest midline under inhalation anesthesia. Adhesive scar tissue was dissected, and the silk thread ligating the aorta was identified by the loop created with 5-0 monofilament thread. The 3-0 silk thread was cut, eliminating the stenosis, taking care to avoid aorta damage. Procedures in the normal heart (NH), negative control (NC), and TAC groups were performed similarly, with the addition of cutting the aortic ligatures in the TAC-release (TAC-R) group. The chest was closed as in the initial surgery, and postoperative pain was managed as described above.

### Echocardiography

Cardiac function was assessed by means of transthoracic echocardiography at 4 and 8 weeks after surgery. Echocardiography was performed with a Lumify L12-4 (Philips, Amsterdam, Netherlands) Linear Array Ultrasound System equipped with a 12- to 4-MHz transducer. The animals were anesthetized with isoflurane gas, and two-dimensional imaging of their hearts was performed, with M-mode recordings acquired through the ventricular septum and posterior LV wall. Interventricular septum and posterior wall thicknesses (both end-diastolic and end-systolic) and LV internal dimensions were measured over at least three consecutive cardiac cycles, according to established protocols^13, 59, 60^. LV volume was determined by M-mode echocardiography; the left ventricular end-diastolic dimension (LVDd) and left ventricular end-systolic dimension (LVDs) were measured on the parasternal short-axis view, assuming the left ventricle to be a spheroid. LV volume was typically calculated according to the Teichholz formula: V = 7.0 × D3 / (2.4 + D)^61^. Left ventricular stroke volume (LVSV) was derived by subtracting left ventricular end-systolic volume (LVESV) from left ventricular end-diastolic volume (LVEDV). LVEF was calculated as follows: LVEF = SV × 100 / LVEDV (%)^61^. Data shown were derived from three independent measurements obtained in a blinded fashion.

### RNA extraction and RT-PCR

Total RNA was extracted from rats’ hearts with use of the Gene Jet PCR Purification Kit (Thermo Fisher Scientific, Waltham, MA) and quantified with a Nano-Drop One spectrophotometer (Thermo Scientific). For this RNA extraction, fresh tissue was excised from the LV wall. cDNA was synthesized from 150 ng of total RNA extracted from heart tissues with use of a High Capacity cDNA Reverse Transcription Kit (Applied Biosystems, Framingham, MA). RT-PCR was performed with QuantStudio (Applied Biosystems) and SYBR Premix Ex Taq II (Takara Bio, Shiga, Japan) reagent under the following conditions: 95°C for 30 seconds followed by 40 cycles at 95°C for 5 seconds and 60°C for 30 seconds. Gene expression levels were normalized to *Gapdh* (forward, 5’-CCCCTGGCCAAGGTCATCCA-3’; reverse, 5’-CGGAAGGCCATGCCAGTGAG-3’) expression. Gene expression data were acquired from independent biological replicates. Primers were *Actc1* (forward, 5’-TTTGGTGTGCGACAATGGCT-3’; reverse, 5’-TGCCCCATACCTACCATGAC-3’), *αSMA* (forward, 5’-ACAACTGGTATTGTGCTGGACT-3’; reverse, 5’-TGGCATGAGGCAGAGCATAG-3’), *Angpt2* (forward, 5’-AGTCAGGACTCACCACCAGT-3’; reverse, 5’-CCTCCACCCATGTCCATGTC-3’), *Col1a1* (forward, 5’-TGAGCCAGCAGATTGAGAAC-3’; reverse, 5’-CCAGTACTCTCCGCTCTTCC-3’), *Col3a1* (forward, 5’-AGTCTGGAGTCGGAGGAATG-3’; reverse, 5’-AGGATGTCCAGAGGAACCAG-3’), *Fgf1* (forward, 5’-CTTACCACAGCAGCAGGAATG-3’; reverse, 5’-AACTGTCGATGGTGCGTTCA-3’), *Fgf2* (forward, 5’-CTCTACTGCAAGAACGGCG-3’; reverse, 5’-ACTCCTCTCTCTTCTGCTTGGA-3’), *Fgf9* (forward, 5’-GGAAAGACCACAGCCGATTC-3’; reverse, 5’-TGATCCATACAGCTCCCCCT-3’), *Fn1* (forward, 5’-CGGAGGCCACCATCACTG-3’; reverse, 5’-GTGGAAGGGTAACCAGTTGGG-3’), *Igf1* (forward, 5’-TACTTCAACAAGCCCACAGG-3’; reverse, 5’-CTCATCCACAATGCCTGTCT-3’), *Igfbp5* (forward, 5’-AACGAAAAGAGCTACGGCGA-3’; reverse, 5’-GACCTTCGGGGAGTAGGTCT-3’), *Il1b* (forward, 5’-TGGCAACTGTCCCTGAACTC-3’; reverse, 5’-AAGGGCTTGGAAGCAATCCTTA-3’), *Mmp2* (forward, 5’-AGAAGGCTGTGTTCTTCGCA-3’; reverse, 5’-AAAGGCAGCGTCTACTTGCT-3’), *Mmp9* (forward, 5’-TGCCCTGGAACTCACACAAC-3’; reverse, 5’-GTTCACCCGGTTGTGGAAAC-3’), *Mybpc3* (forward, 5’-AGGGCATGAAGCACGATGAA-3’; reverse, 5’-AGCCAGTTCCACAGTAAGCC-3’), *Myl2* (forward, 5’-CAGCAGCCAGCCTCAGAC-3’; reverse, 5’-CCGTCTCTGTTCTGGTCCAT-3’), *Myl3* (forward, 5’-AATCCTACCCAGGCAGAGGT-3’; reverse, 5’-TGTGCCCGTGTCTTTGTTCT-3’), *Pdgfb* (forward, 5’-GGAGTCGGCATGAATCGCT-3’; reverse, 5’-ATGGAGTGGTCACTCAGCAT-3’), *Rcan1* (forward, 5’-CTGCCCGTTGAAAAAGCAGAA-3’; reverse, 5’-AATTTGGCCCTGGTCTCACTT-3’), *Tgfb1* (forward, 5’-CAACTTCTGTCTGGGACCCT-3’; reverse, 5’-CGGGTTGTGTTGGTTGTAGA-3’), *Timp1* (forward, 5’-TTTCTGCAACTCGGACCTGG-3’; reverse, 5’-CGTCGAATCCTTTGAGCATCTTA-3’), *TNFα* (forward, 5’-TCGTAGCAAACCACCAAGTG-3’; reverse, 5’-TTGTCTTTGAGATCCATGCC-3’), *Tnni3* (forward, 5’-GGGACACCCTTCTAAGACCC-3’; reverse, 5’-CATCGCTGCTCTCATCCGC-3’), *Tnnt2* (forward, 5’-CTGAGGCTGAGCAGACACC-3’; reverse, 5’-GCGTCAGACATGTTCTCCGT-3’), *Tmp1* (forward, 5’-TCCAGCTGAAAGAGGCCAAG-3’; reverse, 5’-CACATTTGCCTTCCGAGAGC-3’), and *Vegfa* (forward, 5’-GTACCTCCACCATGCCAAGT-3’; reverse, 5’-GCATTCACATCTGCTGTGCT-3’).

### mtDNA copy numbers

Total DNA, including mtDNA, was extracted from heart tissue with use of the ISOSPIN Tissue DNA Kit (NIPPON GENE, Tokyo, Japan) and quantified with a Nano-Drop One spectrophotometer (Thermo Scientific). For DNA extraction from rat hearts, fresh tissue was excised from the left ventricular wall. RT-PCR was performed with use of the QuantStudio (Applied Biosystems) and SYBR Premix Ex Taq II (Takara Bio) under the following conditions: 98℃ for 2 minutes followed by 40 cycles at 98℃ for 10 seconds, 60℃ for 15 seconds, and 68℃ for 30 seconds. RT-PCR was performed on one sample by using four primer pairs, and the mtDNA copy number was calculated by relative quantification based on the difference in cycle threshold values between mtDNA and nuclear DNA. Primers used were *mt-ND1* (forward, 5’-ATCTGTGTTATTAGACATGCACCA-3’; reverse, 5’-TAGTCACCCCCAGGACGAAT-3’), *mt-ND5* (forward, 5’-ACATGCACCATTAAGTCATAAACCT-3’; reverse, 5’-ATAGTCACCCCCAGGACGAA-3’), *SLCO2B1* (forward, 5’-TCAGCCCTTTGGCATCTCC-3’; reverse, 5’-CATCATGGTCACTGCAAACAGG-3’), and *SERPINA1* (forward, 5’-AGCCAGGGAGACAGGGA-3’; reverse, 5’-AGCCAGGGAGACAGGGA-3’).

### Microarray analysis

Total RNA was isolated from whole heart according to the aforementioned procedure and analyzed by means of GeneChip technology (GeneChip Arrays platform with Clariom™ S Assay, Rat; Thermo Fisher Scientific). Standard quality control steps were included to determine total RNA quality with use of Agilent Bioanalyzer 2100 (RNA integrity number (RIN) > 7.0; Agilent Technologies, Santa Clara, CA) and quantity with use of a nanoliter spectrophotometer (Nanodrop, Thermo Fisher Scientific). Two independent biological replicates were prepared from whole hearts. Median per-chip normalization was applied to each array. Differentially expressed genes were identified at a fold-change cutoff of 2.0.

### Immunohistochemistry

For immunohistochemical analysis, rats were euthanized under deep anesthesia with complete respiratory and cardiac arrest. Cervical dislocation was then added to achieve reliable euthanasia, and this was followed immediately by clamping of the aorta and injection of ice-cold PBS into the left ventricular cavity. The heart was then perfused with ice-cold 4% paraformaldehyde in PBS. Subsequently, the heart was excised, bisected at the midpoint along the short axis of the left ventricle, embedded in optimal cutting temperature compound (VWR International, Radnor, PA), and rapidly frozen in isopentane chilled in liquid nitrogen. Frozen tissue sections (8 µm thick) were prepared, and non-specific antibody-binding sites were blocked with PBS containing 5% goat serum. Antibodies were then applied overnight at 4°C: Griffonia Simplicifolia Ⅰ-Isolectin B4, DyLight 594 Conjugate at 1:100 dilution (DL-1207, Vector Laboratories, Burlingame, CA) and anti-Cardiac Troponin T Ab at 1:00 dilution (15513-1-AP, Proteintech, Rosemont, IL). Slides were washed and, if necessary, incubated with fluorophore-conjugated secondary antibody at 1:300 dilution (Alexa Fluor 488-conjugated polyclonal antibody, Invitrogen, Waltham, MA) and DAPI in blocking buffer for 1 hour at room temperature. Data were acquired from independent biological replicates.

### Masson’s Trichrome staining

Frozen tissue sections (8 µm thick) were prepared as described above and stained with trichrome (Trichrome Stain Kit, Scytek Laboratories, Logan, UT) according to the manufacturer’s instructions. Data were acquired from independent biological replicates.

### Imaging and analysis

Digital images were captured with use of an all-in-one fluorescence microscope (BZ-8000; KEYENCE, Osaka, Japan) and imported as TIFF files into ImageJ (National Institutes of Health). Images were obtained from three separate regions, and the fibrin-positive area in each region was quantified as a percentage of the total region area. The Color Threshold function in ImageJ was used to measure the fibrin clot area. Initially, the image was divided into three parts by means of the Split Channels function, and the clearest view of the target area was selected. Subsequently, the threshold was adjusted and set to detect and highlight only the object of interest (i.e., the fibrin clot area). The percentage of the object area in the image was then calculated. This procedure was applied to both Control and target group images, with the threshold value set for the Control group also used for the target group.

### In vivo transfection

To investigate the role of RCAN1 in cardiac fibrosis, in vivo gene silencing was performed by using *Rcan1* siRNA (AM16706, Invitrogen) along with a transfection reagent, according to a previously described protocol^62^ with modifications. Forty micrograms of siRNA, combined with non-viral in vivo transfection reagent JetPEI® (Polyplus-transfection), was administered to each animal three times per week by tail-vein injection. An N/P ratio of 8 in a total volume of 500 μL of 5% dextrose was maintained for each injection, according to the manufacturer’s protocol. NC siRNA (AM4635, Invitrogen) was administered for negative control.

### Statistical analysis

Statistical analysis was performed with GraphPad Prism version 8 (GraphPad Software, San Diego, CA). Data were assumed to be normally distributed and are presented as mean ± SEM. All statistical tests were parametric. For comparison between multiple sample sets, repeated-measures analysis of variance (ANOVA) or one- or two-way ANOVA was performed, followed by Bonferroni’s post-hoc test. To evaluate differences between two sample sets, two-tailed, unpaired Student’s t-test was applied.

## Supporting information

Support Figure1

## Acknowledgements

This project was supported by SENSHIN Medical Research Foundation, a Grant-in-Aid for Scientific Research (C), TERUMO LIFE SCIENCE FOUNDATION, Suzuken Memorial Foundation, Kato Memorial Trust for Nambyo Research, a grant from Bristol Myers Squibb for non-clinical research. The author thanks Ms. Wendy Alexander-Adams for her assistance in reporting the findings in English.

## Author contributions

M.S. conceptualized and designed the study; performed the experiments, acquired, analyzed, and interpreted the data; and drafted and critically revised the manuscript.

## Data availability statement

The author declares that all supporting data are available within the article and its online supplementary file. For purposes of reproducibility, additional technical information and data that support the findings of this study are available from the corresponding author on reasonable request. Microarray data can be obtained from Gene Expression Omnibus GSE270186 https://www.ncbi.nlm.nih.gov/geo/query/acc.cgi?acc=GSE270186

## References

1. Writing Committee, M., et al. 2016 ACC/AHA/HFSA Focused Update on New Pharmacological Therapy for Heart Failure: An Update of the 2013 ACCF/AHA Guideline for the Management of Heart Failure: A Report of the American College of Cardiology/American Heart Association Task Force on Clinical Practice Guidelines and the Heart Failure Society of America. Circulation 134, e282–293 (2016).

2. Rockman, H.A. et al. Segregation of atrial-specific and inducible expression of an atrial natriuretic factor transgene in an in vivo murine model of cardiac hypertrophy. Proc Natl Acad Sci U S A 88, 8277–8281 (1991).

3. You, J. et al. Differential cardiac hypertrophy and signaling pathways in pressure versus volume overload. Am J Physiol Heart Circ Physiol 314, H552–H562 (2018).

4. Bao, D. et al. Tomoregulin-1 prevents cardiac hypertrophy after pressure overload in mice by inhibiting TAK1-JNK pathways. Dis Model Mech 8, 795–804 (2015).

5. Heineke, J. & Molkentin, J.D. Regulation of cardiac hypertrophy by intracellular signalling pathways. Nat Rev Mol Cell Biol 7, 589–600 (2006).

6. Hill, J.A. & Olson, E.N. Cardiac plasticity. N Engl J Med 358, 1370–1380 (2008).

7. Janicki, J.S. & Brower, G.L. The role of myocardial fibrillar collagen in ventricular remodeling and function. J Card Fail 8, S319–325 (2002).

8. Kong, P., Christia, P. & Frangogiannis, N.G. The pathogenesis of cardiac fibrosis. Cell Mol Life Sci 71, 549–574 (2014).

9. Dewald, O. et al. Development of murine ischemic cardiomyopathy is associated with a transient inflammatory reaction and depends on reactive oxygen species. Proc Natl Acad Sci U S A 100, 2700–2705 (2003).

10. Brilla, C.G., Matsubara, L. & Weber, K.T. Advanced hypertensive heart disease in spontaneously hypertensive rats. Lisinopril-mediated regression of myocardial fibrosis. Hypertension 28, 269–275 (1996).

11. Broughton, K.M. et al. Mechanisms of Cardiac Repair and Regeneration. Circ Res 122, 1151–1163 (2018).

12. Dimmeler, S. & Zeiher, A.M. Endothelial cell apoptosis in angiogenesis and vessel regression. Circ Res 87, 434–439 (2000).

13. Shiraishi, M., Suzuki, K. & Yamaguchi, A. Effect of mechanical tension on fibroblast transcriptome profile and regulatory mechanisms of myocardial collagen turnover. FASEB J 37, e22841 (2023).

14. Zhou, Y. et al. Metascape provides a biologist-oriented resource for the analysis of systems-level datasets. Nat Commun 10, 1523 (2019).

15. Yellon, D.M. & Hausenloy, D.J. Myocardial reperfusion injury. N Engl J Med 357, 1121–1135 (2007).

16. Chouchani, E.T. et al. A Unifying Mechanism for Mitochondrial Superoxide Production during Ischemia-Reperfusion Injury. Cell Metab 23, 254–263 (2016).

17. Hausenloy, D.J. & Yellon, D.M. Myocardial ischemia-reperfusion injury: a neglected therapeutic target. J Clin Invest 123, 92–100 (2013).

18. Duan, H. et al. Rcan1-1L overexpression induces mitochondrial autophagy and improves cell survival in angiotensin II-exposed cardiomyocytes. Exp Cell Res 335, 99–106 (2015).

19. Norberg, K.J. et al. RCAN1 is a marker of oxidative stress, induced in acute pancreatitis. Pancreatology 18, 734–741 (2018).

20. Wang, S., Wang, Y., Qiu, K., Zhu, J. & Wu, Y. RCAN1 in cardiovascular diseases: molecular mechanisms and a potential therapeutic target. Mol Med 26, 118 (2020).

21. Chen, E.Y. et al. Enrichr: interactive and collaborative HTML5 gene list enrichment analysis tool. BMC Bioinformatics 14, 128 (2013).

22. Dellsperger, K.C. & Marcus, M.L. The effects of pressure-induced cardiac hypertrophy on the functional capacity of the coronary circulation. Am J Hypertens 1, 200–207 (1988).

23. Eltzschig, H.K. & Eckle, T. Ischemia and reperfusion--from mechanism to translation. Nat Med 17, 1391–1401 (2011).

24. Brandes, R.P., Weissmann, N. & Schroder, K. Redox-mediated signal transduction by cardiovascular Nox NADPH oxidases. J Mol Cell Cardiol 73, 70–79 (2014).

25. Tsutsui, H., Kinugawa, S. & Matsushima, S. Mitochondrial oxidative stress and dysfunction in myocardial remodelling. Cardiovasc Res 81, 449–456 (2009).

26. Murphy, E. & Steenbergen, C. Mechanisms underlying acute protection from cardiac ischemia-reperfusion injury. Physiol Rev 88, 581–609 (2008).

27. Molkentin, J.D. et al. A calcineurin-dependent transcriptional pathway for cardiac hypertrophy. Cell 93, 215–228 (1998).

28. Duerr, R.L. et al. Insulin-like growth factor-1 enhances ventricular hypertrophy and function during the onset of experimental cardiac failure. J Clin Invest 95, 619–627 (1995).

29. Page, S.P. et al. Cardiac myosin binding protein-C mutations in families with hypertrophic cardiomyopathy: disease expression in relation to age, gender, and long term outcome. Circ Cardiovasc Genet 5, 156–166 (2012).

30. Kimura, A. et al. Mutations in the cardiac troponin I gene associated with hypertrophic cardiomyopathy. Nat Genet 16, 379–382 (1997).

31. Watkins, H. et al. Mutations in the genes for cardiac troponin T and alpha-tropomyosin in hypertrophic cardiomyopathy. N Engl J Med 332, 1058–1064 (1995).

32. Mogensen, J. et al. Alpha-cardiac actin is a novel disease gene in familial hypertrophic cardiomyopathy. J Clin Invest 103, R39–43 (1999).

33. Mavilakandy, A. & Ahamed, H. Mutation of the MYL3 gene in a patient with mid-ventricular obstructive hypertrophic cardiomyopathy. BMJ Case Rep 15 (2022).

34. Flavigny, J. et al. Identification of two novel mutations in the ventricular regulatory myosin light chain gene (MYL2) associated with familial and classical forms of hypertrophic cardiomyopathy. J Mol Med (Berl) 76, 208–214 (1998).

35. Alfieri, C.M., Evans-Anderson, H.J. & Yutzey, K.E. Developmental regulation of the mouse IGF-I exon 1 promoter region by calcineurin activation of NFAT in skeletal muscle. Am J Physiol Cell Physiol 292, C1887–1894 (2007).

36. Knoll, R. Myosin binding protein C: implications for signal-transduction. J Muscle Res Cell Motil 33, 31–42 (2012).

37. Wang, M. et al. TRPA1 deficiency attenuates cardiac fibrosis via regulating GRK5/NFAT signaling in diabetic rats. Biochem Pharmacol 214, 115671 (2023).

38. Saygili, E. et al. The angiotensin-calcineurin-NFAT pathway mediates stretch-induced up-regulation of matrix metalloproteinases-2/-9 in atrial myocytes. Basic Res Cardiol 104, 435–448 (2009).

39. Deschamps, A.M. & Spinale, F.G. Pathways of matrix metalloproteinase induction in heart failure: bioactive molecules and transcriptional regulation. Cardiovasc Res 69, 666–676 (2006).

40. Fan, D., Takawale, A., Lee, J. & Kassiri, Z. Cardiac fibroblasts, fibrosis and extracellular matrix remodeling in heart disease. Fibrogenesis Tissue Repair 5, 15 (2012).

41. Eghbali, M. Cardiac fibroblasts: function, regulation of gene expression, and phenotypic modulation. Basic Res Cardiol 87 **Suppl 2**, 183–189 (1992).

42. Butt, R.P., Laurent, G.J. & Bishop, J.E. Collagen production and replication by cardiac fibroblasts is enhanced in response to diverse classes of growth factors. Eur J Cell Biol 68, 330–335 (1995).

43. Matsusaka, H. et al. Targeted deletion of matrix metalloproteinase 2 ameliorates myocardial remodeling in mice with chronic pressure overload. Hypertension 47, 711–717 (2006).

44. Heymans, S. et al. Loss or inhibition of uPA or MMP-9 attenuates LV remodeling and dysfunction after acute pressure overload in mice. Am J Pathol 166, 15–25 (2005).

45. Lu, J. et al. Amlodipine and Atorvastatin Improved Hypertensive Cardiac Remodeling through Regulation of MMPs/TIMPs in SHR Rats. Cell Physiol Biochem 39, 47–60 (2016).

46. Varo, N. et al. Chronic AT(1) blockade stimulates extracellular collagen type I degradation and reverses myocardial fibrosis in spontaneously hypertensive rats. Hypertension 35, 1197–1202 (2000).

47. Yang, L. et al. VEGF increases the proliferative capacity and eNOS/NO levels of endothelial progenitor cells through the calcineurin/NFAT signalling pathway. Cell Biol Int 36, 21–27 (2012).

48. Manabe, T., Park, H. & Minami, T. Calcineurin-nuclear factor for activated T cells (NFAT) signaling in pathophysiology of wound healing. Inflamm Regen 41, 26 (2021).

49. Jabr, R.I. et al. Nuclear translocation of calcineurin Abeta but not calcineurin Aalpha by platelet-derived growth factor in rat aortic smooth muscle. Am J Physiol Cell Physiol 292, C2213–2225 (2007).

50. Minami, T. et al. The calcineurin-NFAT-angiopoietin-2 signaling axis in lung endothelium is critical for the establishment of lung metastases. Cell Rep 4, 709–723 (2013).

51. Oka, T., Akazawa, H., Naito, A.T. & Komuro, I. Angiogenesis and cardiac hypertrophy: maintenance of cardiac function and causative roles in heart failure. Circ Res 114, 565–571 (2014).

52. Kutryk, M.J. & Stewart, D.J. Angiogenesis of the heart. Microsc Res Tech 60, 138–158 (2003).

53. Carmeliet, P. & Jain, R.K. Molecular mechanisms and clinical applications of angiogenesis. Nature 473, 298–307 (2011).

54. Varricchi, G., et al. Angiopoietins, Vascular Endothelial Growth Factors and Secretory Phospholipase A(2) in Ischemic and Non-Ischemic Heart Failure. J Clin Med 9 (2020).

55. Rotter, D., et al. Calcineurin and its regulator, RCAN1, confer time-of-day changes in susceptibility of the heart to ischemia/reperfusion. J Mol Cell Cardiol 74, 103–111 (2014).

56. deAlmeida, A.C., van Oort, R.J. Wehrens, X.H. Transverse aortic constriction in mice. J Vis Exp (2010).

57. Zaw, A.M., Williams, C.M., Law, H.K. & Chow, B.K. Minimally Invasive Transverse Aortic Constriction in Mice. J Vis Exp (2017).

58. Liu, B., Li, A., Gao, M., Qin, Y. & Gong, G. Modified Protocol for A Mouse Heart Failure Model Using Minimally Invasive Transverse Aortic Constriction. STAR Protoc 1, 100186 (2020).

59. Shiraishi, M., Yamaguchi, A. & Suzuki, K. Nrg1/ErbB signaling-mediated regulation of fibrosis after myocardial infarction. FASEB J 36, e22150 (2022).

60. Shiraishi, M. et al. Alternatively activated macrophages determine repair of the infarcted adult murine heart. J Clin Invest 126, 2151–2166 (2016).

61. Shiraishi, M., Kimura, N. & Yamaguchi, A. Early cardiac contractility outcome of reoperative coronary artery bypass grafting using right gastroepiploic artery. J Card Surg 36, 4103–4110 (2021).

62. Garg, M. et al. Cardiolipin-mediated PPARgamma S112 phosphorylation impairs IL-10 production and inflammation resolution during bacterial pneumonia. Cell Rep 34, 108736 (2021).

