## Supplementary material for "Unveiling reversibility and plasticity in cardiac hypertrophy: insights from a transverse aortic constriction-release model": Support Figure1

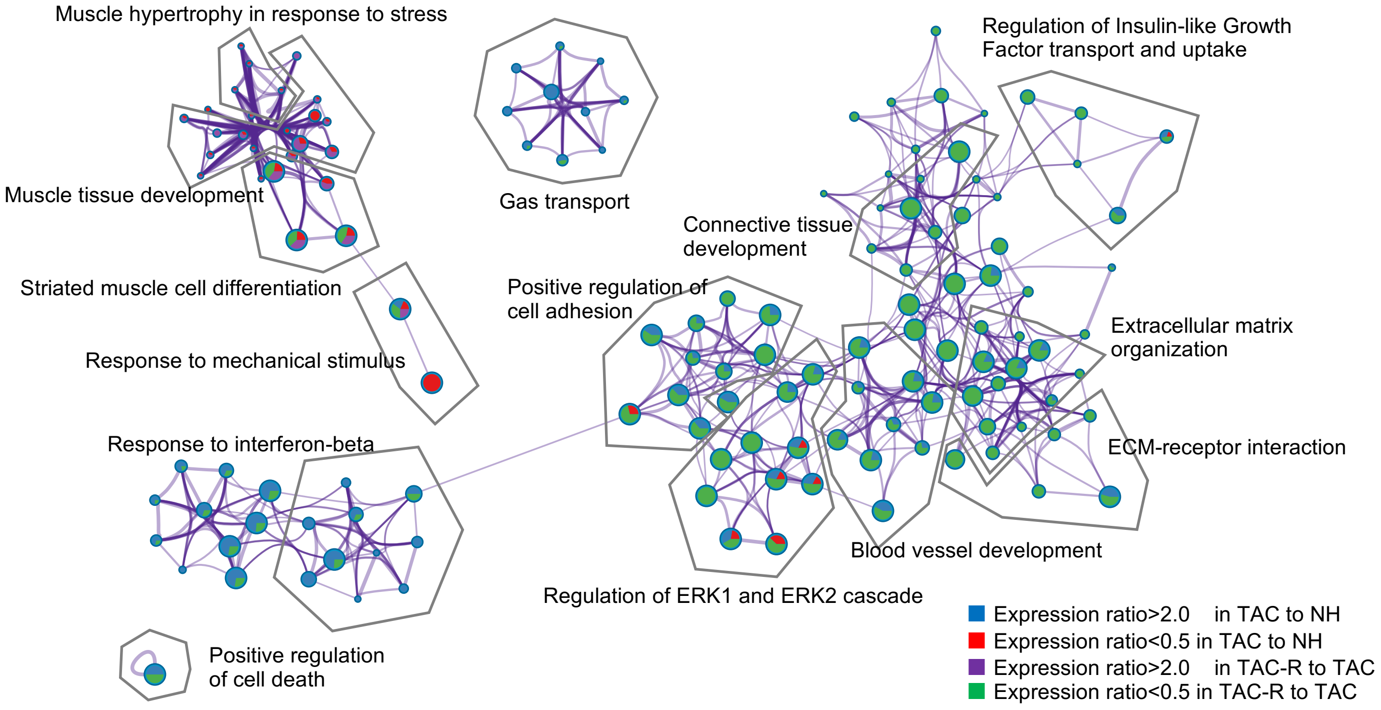


**Supplementary Figure 1. Subset of representative terms and network layout.**
Each term is represented by a circular node, with the size of each node proportional to the number of input genes that fall under that term and the color of each node indicative of its cluster identity. Terms with a similarity score > 0.3 are linked. Thickness of the edge represents the similarity score. The Cytoscape layout is force-directed. Each term label serves as an aggregate of the labels in each cluster.
